# Synergistic olfactory processing for social plasticity in desert locusts

**DOI:** 10.1101/2023.09.15.557953

**Authors:** Inga Petelski, Yannick Günzel, Sercan Sayin, Susanne Kraus, Einat Couzin-Fuchs

**Affiliations:** International Max Planck Research School for Quantitative Behavior, _Ecology and Evolution from lab to field_, _78464 Konstanz_, Germany; Department of Biology, University of Konstanz, _78464 Konstanz_, Germany; Department of Collective Behavior, Max Planck Institute of Animal Behavior, _78464 Konstanz_, Germany; Centre for the Advanced Study of Collective Behaviour, University of Konstanz, _78464 Konstanz_, Germany

**Keywords:** Locusts, Olfaction, Calcium imaging, Social modulation

## Abstract

1

Desert locust plagues threaten the food security of millions. Central to their formation is crowding-induced social plasticity from ‘solitarious’ to ‘gregarious‘ phenotypes. We investigated the impact of population density changes on locusts’ foraging choices and their neurobiology by examining how relevant food and social odors are coded in the antennal lobe. Our analysis shows that gregarious locusts are highly attentive to social cues during foraging, with olfaction playing an essential role. Using calcium imaging, we show that corresponding odors are encoded by projection neurons, revealing a stable combinatorial response map. Transient dynamics in the glomeruli converge into temporally evolving response motifs in the somata that differ between gregarious and solitarious insects. The dynamics of response motifs facilitate a crowding-dependent synergy between olfactory processing of food-related and social odors. Our study demonstrates the effectiveness of calcium imaging for locust olfaction, suggesting a crowding-induced adaptation to enhance food detection in swarms.

**Teaser:** In dense swarms, desert locusts optimize foraging efficiency by exhibiting an enhanced olfactory response to food odors.

## 3 Introduction

The ability to adapt to a changing social environment is a fundamental aspect of life. Locusts exhibit the distinct ability to switch between a solitary and gregarious lifestyle rapidly, rendering them an ideal model for studying social plasticity and the emergence of collective behavior. Furthermore, these transitions are central to the formation of large-scale destructive locust outbreaks, which are precipitated by an autocatalytic process of positive feedback between population density and aggregation behavior (Cullen et al. (2017); Pener and Simpson (2009); Simpson et al. (2009, 1999)). When population density spontaneously increases, formerly solitarious locusts are forced to conglomerate on limited resources, leading to further increases in local density and subsequent directed aggregation of animals that have become gregarious. These dynamics also trigger changes in other density-dependent traits, including metabolism, developmental, and reproductive physiology, which together foster population growth and swarm formation (cumulatively termed ’phase change’, (Sword et al. (2010))). As crowding further increases, the demand for nutrients rises, compelling large locust aggregations to leave areas with low food availability forage despite associated risks of predation and cannibalism (Chang et al. (2023); Collett et al. (1998); Guttal et al. (2012); Sword et al. (2000); Wei et al. (2019)). These rapid density-dependent adaptations pose questions about the neural processing in changing social environments that allow animals to adjust their decision-making appropriately in a context-dependent manner.

The transition to group living also alters the sensory information available to locusts when searching and selecting feeding sites. Gregarious desert locusts (*Schistocerca gregaria*) effectively utilize both asocial (*e.g.*, sight and/or smell of food) and social (*e.g.*, presence or action of conspecifics) cues for their foraging decisions (Günzel et al. (2023)). Specifically, the decision to join a food patch is positively influenced by the number of conspecifics on the patch, as well as by its quality and the locust’s prior experience with it. However, our understanding of the sensory processes that mediate these density-dependent decisions remains limited.

Like many animals, locusts use olfactory information to locate food, find mates, and avoid predators or toxins. Invariably, olfactory systems face the challenge of encoding and decoding a vast array of chemical structures across varying odor concentrations and mixtures in turbulent environments. Knowledge gained about olfactory systems across the animal kingdom has demonstrated a largely typical organization of a single neuropile — the antennal lobe (AL) in insects and the olfactory bulb in mammals — located only one synapse away from the peripheral sensory neurons (Wilson and Mainen (2006)). To meet the challenge of complexity in chemical space, the antennal lobe (like the olfactory bulb) acts as an early processing center in which numerous incoming olfactory receptor neurons (ORNs) converge on fewer output projection neurons (PNs). Non-linear summation in such synapses and gain control mediated by local neurons enables the coding of odors over dynamic ranges (Galizia (2014)).

Invertebrate and vertebrate model systems in olfactory research, including rodents (Stewart et al. (1979)), zebrafish (Korsching et al. (1999)), and fruit flies (Rodrigues (1988)), have a typical early olfactory coding architecture. Convergent input and output neurons form a computational unit called the glomerulus. Each glomerulus receives input from all ORNs expressing the same olfactory receptor type, and most output neurons are exclusive to a particular glomerulus (Wilson and Mainen (2006)). Such a glomerular structure effectively organizes olfactory coding in linear and parallel channels. In locusts, however, the AL structure and connectivity deviate from those of many insects (Ignell et al. (2001)). In contrast to, for example, 50 individually identified glomeruli in the fruit fly (Hallem and Carlson (2004)), it houses an unusually large number of more than a thousand so-called microglomeruli and more strikingly, all output PNs innervate multiple microglomeruli (no uniglomerular PNs have been described to date), and each ORN projects to multiple glomeruli (Anton and Hansson (1996); Hansson et al. (1996); Hansson and Stensmyr (2011); Ignell et al. (1998, 2001)). Consequently, it is largely unknown how this structural difference impacts locust olfaction and if it is related to the notable phenotypic plasticity many locust species exhibit.

While the wiring of the locust antennal lobe remains unresolved, research on odor processing in locusts has played a pioneering role in advancing our understanding of combinatorial coding in olfactory systems (Laurent and Davidowitz (1994); Laurent et al. (1996); Wehr and Laurent (1996)). Using intracellular and extracellular recording techniques and computational models, locust odor representation has been postulated as a dynamic combinatorial code (Anton and Hansson (1996); Ignell et al. (1998)) suitable for rapid discrimination of odor identity and intensity (Mazor and Laurent (2005); Stopfer et al. (2003)), as well as for contrast enhancement when stimuli overlap or follow distractor odors (Broome et al. (2006); Brown et al. (2005); Nizampatnam et al. (2018)). Despite the importance of combinatorial coding and ensemble dynamics, most data sets captured only a fraction of antennal lobe output at a given recording or lacked the cell’s identity, which prevented assigning structure to function. Both problems can be addressed with functional imaging techniques that allow monitoring of network-spanning dynamics, which is particularly important for the highly distributed innervation of the locust AL.

To this end, we have established a functional imaging protocol to investigate the spatial organization of odor-evoked activity in nearly all PNs of the locust AL. We map the interaction between food and social odors in gregarious and solitarious locusts to examine how odor representation is impacted by crowding. We link our findings to the animals’ preferences in a behavioral assay by estimating the contribution of different sensory cues to foraging decisions. Our results reveal a crowding-induced olfactory modulation and advance the accessibility of locusts as a model system for studying collective behavior and the neuronal mechanisms underlying social plasticity.

## 4 Results

### 4.1 Social plasticity and the sensory basis of foraging decisions

To start investigating how population density is linked with sensory processing and decision-making, we analyzed the contribution of different sensory modalities to foraging decisions, with a focus on social and food olfactory cues. Tests were made in simple patch selection assays that presented locusts with four options: food (blackberry leaves, denoted as *Lvs*), a social setting (eight gregarious locusts, *Lct*), a combination of leaves and locusts (*LvsLct*), and an empty control patch (*Ctr*). We observed that gregarious desert locusts, *Schistocerca gregaria*, exhibited a strong attraction to the patch containing both leaves and locusts, as shown in Fig. 1Ai (*Vis*^(+)^*Ol f* ^(+)^). This attraction diminished in the absence of either visual (Fig. 1Aii; *Vis*^(^*^−^*^)^*Ol f* ^(+)^) or olfactory cues (Fig. 1Aiii; *Vis*^(+)^*Ol f* ^(^*^−^*^)^, see also suppl Fig. 1 for animal trajectories). In contrast, solitarious locusts typically disregarded social stimuli and predominantly approached patches where food was presented alone (Fig. 1C). Notably, just as with gregarious animals, the choices of solitarious locusts were most pronounced when both visual and olfactory cues were available.

**Figure 1:**
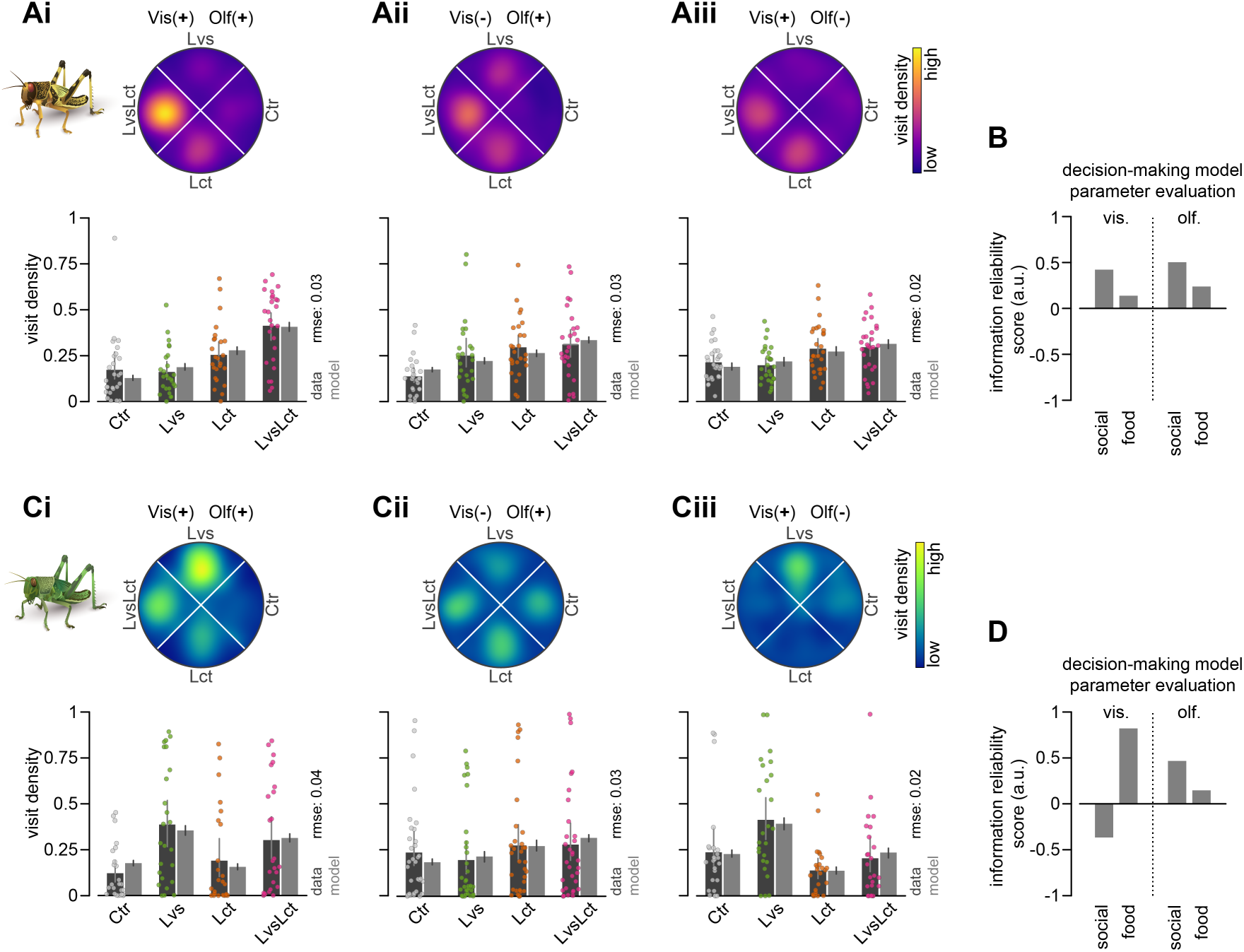
Interactions between social and food cues in patch selection. (A) Gregarious locusts were tested in a circular arena with four patch options under three sensory conditions, designed to isolate the effects of visual and olfactory cues during patch selection. Heatmaps depict average locust density in each stimulus quarter (blackberry leaves *Lvs*, locusts *Lct*, leaves and locusts *LvsLct*, and control *Ctr*) for different sensory conditions (visual and olfactory cues available: *Vis*^(+)^*Ol f* ^(+)^, Ai; olfactory but no visual cues: *Vis*^(^*^−^*^)^*Ol f* ^(+)^, Aii; visual but no olfactory cues: *Vis*^(+)^*Ol f* ^(^*^−^*^)^, Aiii). Swarm plots represent individual locusts’ mean density in each stimulus quarter, dark gray bar plots with error bars indicate corresponding grand means with 95% confidence intervals, and light gray bar plots with error bars indicate means and 95% credible intervals from a decision-making model (5000 simulations; see Methods). (B) Information reliability scores for each information class (social, food) and sensory modality (visual, olfactory), obtained by fitting a decision-making model to the patch selection data in (A). (C-D) Similar experiments and analysis as in A-B, but for solitarious animals.

To dissect the contributions of different sensory cues to the decision process, we extended on a Bayesian decision-making model (Arganda et al. (2012); Günzel et al. (2023)) to encompass the four information classes available in the current assay: vision, olfaction, social, and food. Specifically, we estimated the reliability of each information class—socio-visual, food-visual, socio-olfactory, and food-olfactory—in order to elucidate how these factors influence locusts’ choices (Fig. 1B&D, light gray bars; reliability parameter *_C_i* in Eqn. 1). The close alignment between our model’s predictions and the observed choices (light and dark gray bars respectively in Fig. 1A&C) suggests that locust decisions can be effectively described by the integration of available sensory cues. This model thereby enabled us to further dissect and quantify the parameters that contributed to these decisions. Notably, we found that gregarious animals were highly responsive to social cues, with the highest information reliability scores associated with the presence of conspecific odors at a patch (Fig. 1B). In contrast, solitarious-reared animals appeared deterred by the sight of conspecifics, as indicated by their negative scores in this context (Fig. 1D).

### 4.2 Functional imaging revealed differences in olfactory processing in gregarious and solitarious locusts

Building on the importance of olfaction in patch discrimination, we continued with mapping how the respective odor cues are represented in the locust antennal lobe (AL). An anatomical reconstruction of the AL, using back-fill labeling both olfactory receptor neurons (ORNs) and projection neurons (PNs), shows the arrangement of PN somata on the anterior surface of the AL, with their dendrites situated beneath, encircling a central fiber core. Here, they form functional units, microglomeruli, with ORN axon terminals (Fig. 2A-C). To monitor local calcium signals as a proxy for neuronal activity, we established a protocol for functional imaging via mass retrograde labeling of the entire population of PNs with the fluorescent calcium indicator Cal-520 (Lock et al. (2015)). This technique was optimized to label the vast majority of PNs, facilitating subsequent optical recordings (see Methods for further details). Furthermore, we developed a data-driven analysis pipeline (Fig. 2D) that segments the highly arborized structure of the locust AL into distinct activity granules and overcomes the limitations of biased manual selection of specific regions of interest (see Fig. 2E&G for example response maps).

**Figure 2:**
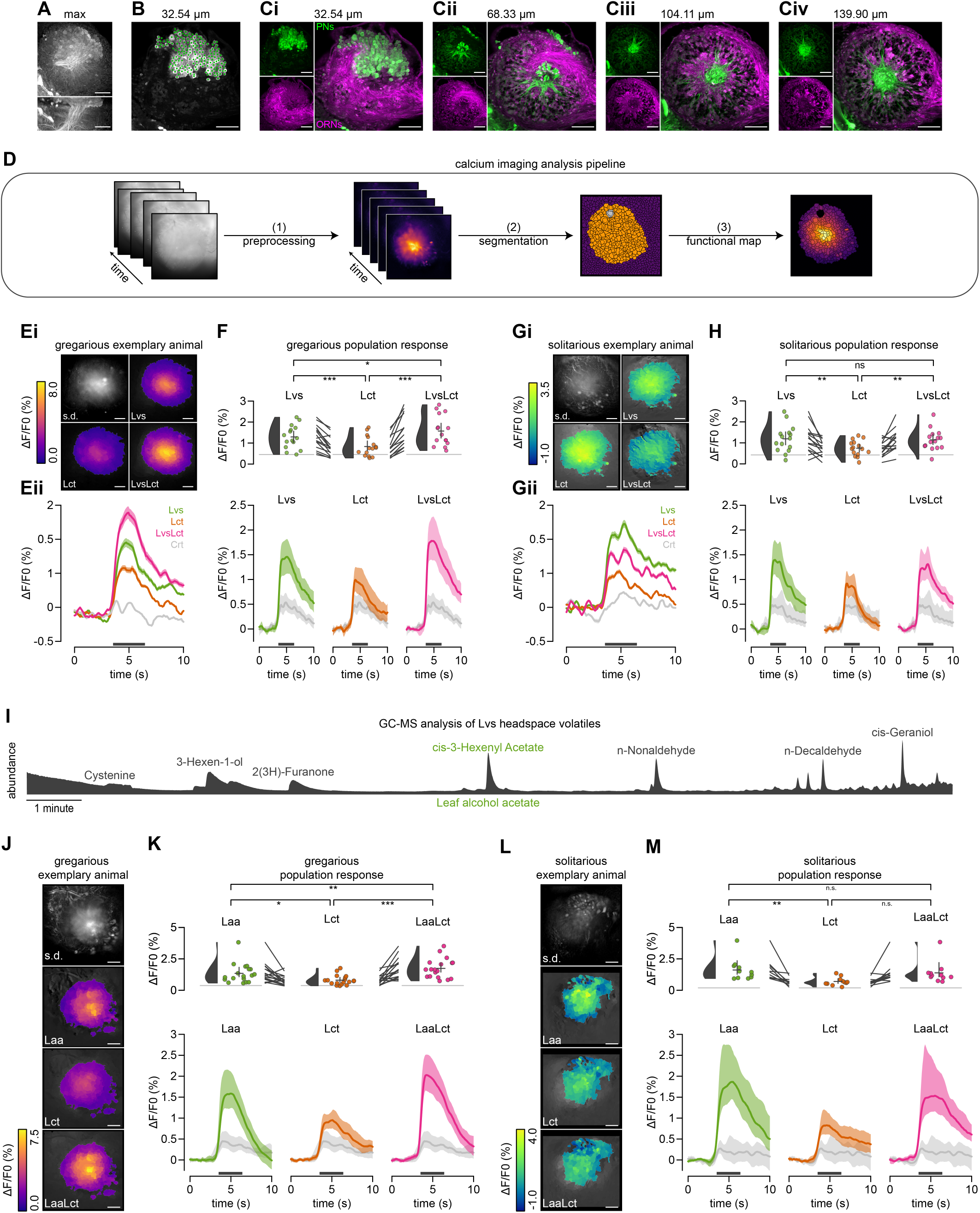
Functional imaging of the antennal lobe representation of the odor cues used in patch choice experiments. (A) Maximum projection of a confocal stack of a projection neuron (PN) backfill staining along the z-axis (top) or the y-axis (bottom). (B) Identification of 181 individually stained PN somata (superimposed green dots) in one section of the scanned locust antennal lobe. (C) Optimal confocal scans with a double backfill staining of PNs (green) and olfactory receptor neurons (ORNs, magenta) at various imaging depths (indicated above, Z=0 corresponds to the anterior surface of the AL). (D) Schematic diagram detailing the staining protocol for PNs in locust antennal lobes for in-vivo functional calcium imaging, and the subsequent analysis pipeline (see Methods for specifics). (E) Widefield calcium imaging data following the presentation of the olfactory stimuli used in the patch selection assay (blackberry leave extract: *Lvs*, locust odor *Lct*, and their combination *LvsLct*). Standard deviation projection across all stimuli (grayscale) of an exemplary gregarious animal, alongside false color-coded images after olfactory stimulation (Ei). Corresponding activity time courses (Eii; mean across all active granules with shaded 95% confidence intervals) for each stimulus (air control in light gray). The dark gray bar specifies the time window for mean response calculations. (F) Average activity time courses of all gregarious animals (grand means with shaded 95% confidence intervals, bottom panels) and respective swarm plots of the individual animal means during the active period (dark gray bars) above, with half-violin plots for probability density estimates, and crosses for grand means and 95% confidence intervals. (G-H) Corresponding analysis as in E-F, but for solitarious animals. (I) Gas chromatography analysis identifies leaf alcohol acetate (*Laa*, cis-3-Hexenyl Acetate) as a dominant volatile in blackberry leaves. (J-M) Correspond to E-H, but for PN responses to the leaf alcohol acetate *Laa* (instead of *Lvs*), the locust odor (*Lct*), and their mixture (*LaaLct*). Statistical inferences in F, H, K, and M were based on paired two-tailed bootstrap randomization tests (n.s.: not significant; p < 0.05: *; p < 0.01: **; p < 0.001: ***), with *p*-value adjustments using the Bonferroni method. Scale bars: 100 *µm*.

Initially, our analysis revealed no significant differences in PN response magnitudes between gregarious and solitarious locusts (N = 15 each) to the leaf odor *Lvs* (*p* = 0.68) or the locust odor *Lct* (*p* = 0.60, two-sample two-tailed bootstrap randomization tests), when these odors were presented separately (Fig. 2F&H showing population response averages for each stimulus with individual data points above). In both, overall PN response to *Lvs* was stronger than to *Lct* (*p <*0.001 and *p <*0.01 for gregarious and solitarious, paired two-tailed bootstrap randomization tests). The comparable response to *Lct* was particularly surprising given that solitarious locusts had minimal, if any, prior exposure to the smell of locusts before the experiment. However, when adding the locust odor to be delivered simultaneously with the leaf odor (*LvsLct*; containing half the amount of each single component — see Methods for mixture preparation), we observed a substantial density-dependent divergence: PN responses of gregarious animals significantly increased (*p* = 0.02, Fig. 2F) upon the addition of the social cue, an increase that was markedly absent in solitarious animals (*p >*0.99, paired two-tailed bootstrap randomization tests; Fig. 2H). These findings suggest that the neural processing of odor mixtures in locusts may be subjected to a density-dependent modulation, highlighting a potential neural mechanism underlying their social behavior.

### 4.3 Response characterization of food and social odors

To further probe the neural underpinnings of the difference in mixture-response, we tested gregarious and solitarious locusts with a related set of single-component odorants and their combinations. Initially, we identified potential food-related candidates from the headspace volatiles of blackberry leaves through gas chromatography-mass spectroscopic analysis. This analysis prominently featured leaf alcohol acetate (cis-3-Hexenyl-acetate, denoted as *Laa*; Fig. 2I), an appetitive plant volatile (Anton and Hansson (1996)). Next, we analyzed the olfactory representation of leaf alcohol acetate *Laa*, the locust odor *Lct*, and their mixture (*LaaLct*), seeking to validate our prior observations of a density-dependent modulation in PN activity (Fig. 2J-M). Similarly as with blackberry leaves, combining *Lct* with *Laa* significantly increased PN activity in gregarious locusts (*p <*0.01, N = 18, Fig. 2K). In solitarious animals, this response was not mirrored (although we note the effect had the same sign as for the gregarious locusts, but was much weaker, *p* = 0.07, N = 11, Fig. 2M). In both cases, the food-related stimulus alone consistently elicited a stronger response than the social cue (*p* = 0.02 for gregarious animals, *p <*0.01 for solitarious animals, paired two-tailed bootstrap randomization tests), but only in gregarious animals the response to the components, when presented together, was stronger than to the individual ones.

To test whether the increased mixture response may relate to a general change in mixture processing, we continued by testing PN responses to two additional mixtures of *Laa* with another, non-locust component: combining equal amounts of *Laa* with 1-Hexanol (*LaaHex*) and with 1-Octanol (*LaaOct*). For gregarious locusts (Fig. 3Ai, same animals as in Figure 2), neither addition—*Hex* nor *Oct*—induced a significant change in PN responses (*p* = 0.71 and *p >*0.99, respectively, paired two-tailed bootstrap randomization tests). In solitarious locusts (Fig. 3Aii), the addition of *Hex* elicited even a significant decrease in comparison with *Laa* alone (*p <*0.01), while *Oct* left responses unchanged (*p* = 0.68). Taken together, these findings suggest that the increased response observed in gregarious locusts when combining the food and social odor, *LaaLct* and *LvsLct*, may represent a specific modulation that heightens the salience of food odors when accompanied by the presence of conspecifics.

**Figure 3:**
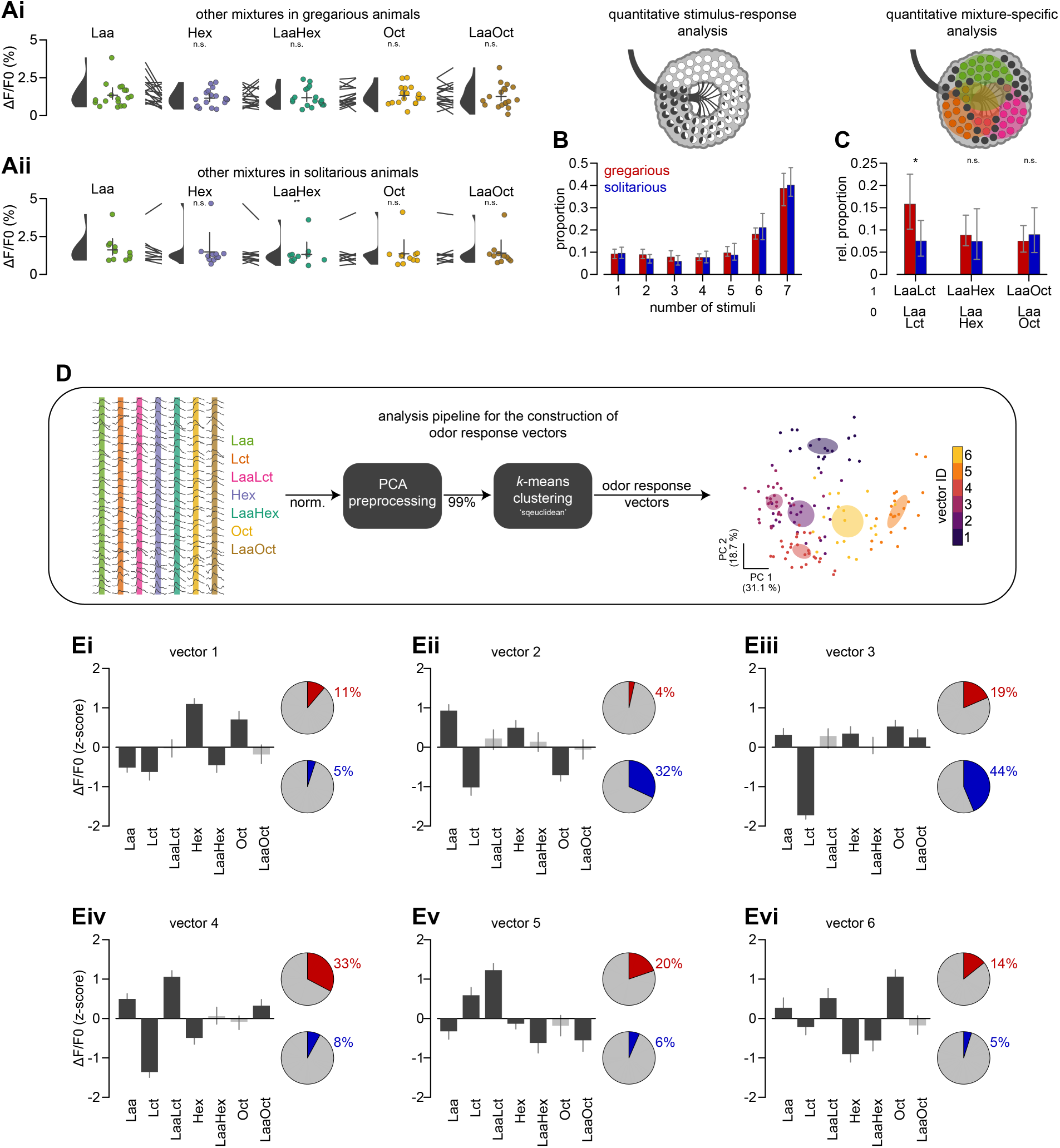
Combinatorial coding of odor responses in antennal lobe projection neurons of gregarious and solitarious animals. (A) Widefield calcium imaging analysis of leaf alcohol acetate (*Laa*, same data as in Fig. 2K&M) and two additional odorants (1-Hexanol *Hex*, 1-Octanol *Oct*), as well as their respective mixtures with *Laa* (*LaaHex*, *LaaOct*) for gregarious (Ai) and solitarious (Aii) animals. (B) The proportion of gregarious (red) and solitarious (blue) granules that respond to a given number of odor stimuli, as symbolized by the pie charts with the proportion of responses for each granule overlaid onto the antennal lobe illustration above. All animals were tested with 7 stimuli: *Laa*, *Lct*, *LaaLct*, *Hex*, *LaaHex Oct*, *LaaOct*. (C) The relative proportion of mixture-specific granules — regions that are responsive to a two-component mixture (1) but not to its two components (0), as symbolized by the Venn diagram overlaid onto the antennal lobe illustration (D) Diagram explaining the clustering of individual granules’ response sets into odor response vectors. The scatter plot displays the first two principal components, colored by cluster identity. Shaded areas mark 99% confidence ellipses, derived from bootstrap resampling (5000 samples). (E) Representation of each odor response vector along with pie charts displaying the relative allocation of gregarious (red) and solitarious (blue) granules to the respective vector. Bars indicate grand means, color-coded to signify if zero is within (light gray) or outside (dark gray) the 95% confidence intervals (gray error bars). Statistical inference in A was based on paired two-tailed bootstrap randomization tests against *Laa*, with *p*-value adjustments using the Bonferroni method. Statistical inference in C was based on unpaired tests. For both n.s.: not significant; p < 0.05: *; p < 0.01: **; p < 0.001: ***.

### 4.4 Combinatorial neural processing with mixture-specific regions in the locust antennal lobe

To better understand how the *LaaLct* mixture is processed and what may mediate the gregarious increase in response magnitude, we further analyzed the odor-induced maps. At first, we noted that despite the considerable overlap in responses to the different stimuli (almost 40% of the granules were active in all seven stimuli, Fig. 3B), the proportion of regions activated by the *LaaLct* mixture but not by its components (mixture-specific regions) was significantly higher in gregarious compared to solitarious animals (Fig. 3C, bars on the left show proportion of *LaaLct*-specific regions that were active for *LaaLct* but non-active for either component, with a similar analysis for the other tested mixtures plotted beside; *p_LaaLct only_* = 0.04, *p_LaaHex only_* = 0.69, *p_LaaOct only_* = 0.63, two-sample two-tailed bootstrap randomization tests).

We projected the response maps from all tested odors and animals into a unified odor response space, as detailed in Fig. 3D. Despite the considerable overlap described above, we discerned distinct odor response vector classes. These classes aided in delineating response types and highlighting potential differences between the phenotypes (Fig. 3E, and Fig. 3D for the visualization of the first two principal components). Upon evaluating these response vectors, we found that certain vectors are primarily driven by the population response to a specific odor. For instance, vector 1 is predominantly influenced by *Hex*, and similarly vector 6 by *Oct*. Conversely, some vectors encapsulate a more combinatorial response profile. For example, vector 3 demonstrates a diverse reaction, with weak positive z-scores for a broad set of stimuli and a strongly negative score for *Lct*. This response type was more common in solitarious locusts (pie charts in Fig. 3E show the proportion of regions for each vector type in red and blue for gregarious and solitarious locusts, respectively). Vectors 4 and 5, on the other hand, were much more prevalent in gregarious animals and are linked to regions with weak sensitivity to either *Laa* or *Lct* but heightened activation upon the introduction of the second mixture component.

### 4.5 Combinatorial odor maps stay consistent across multiple repetitions

Previous studies on locust odor mapping have shown that although the PN population response reliably determines odor identity, there can be substantial variability at the individual PN level (Mazor and Laurent (2005)). We continued our investigation with functional confocal microscopy that allows for detailed monitoring of the spatial arrangement of odor-evoked activities, starting with a test of response consistency in PN somata and glomeruli. In Fig. 4A, we provide a comprehensive visualization of the spatial representation of the general odorants used in our study (*Laa*, *Hex*, and *Oct*). Each stimulus elicited a response in a large set of PNs, apparent as calcium signals both in the somata (Fig. 4Ai, activity granules that match in size, shape, and numbers to individual cell bodies) and concentrically arranged glomeruli (Fig. 4Aii). The overall count of PN cells aligned well with the anatomical reconstruction (Fig. 2B), with the recording planes chosen to maximize the number of detectable somata and glomeruli. As expected from a combinatorial coding system, there was considerable overlap in odor-induced responses (color-coded by odor and displayed as an overlay in Fig. 4Ai & Aii). However, these responses were distinct across different odors and remained consistent for the same odor over multiple trials (see Fig. 4B and suppl. Fig. 2 for individual trials in the example animal, and Fig. 4C for the population response; N = 11 gregarious locusts).

**Figure 4:**
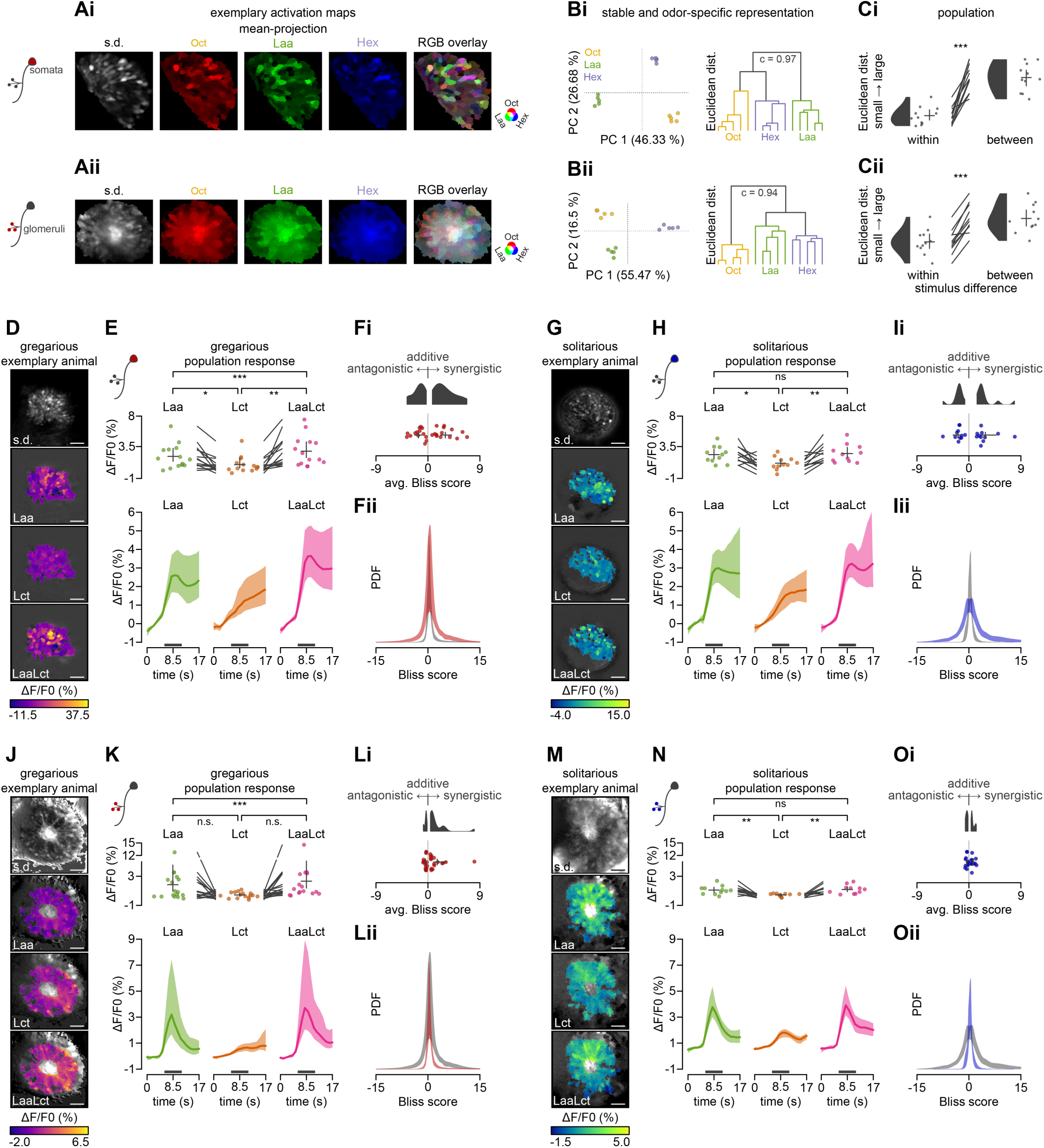
Functional confocal laser scanning microscopy analysis of olfactory processing in somata and glomeruli. (A) Examining the consistency of projection neuron (PN) somata (Ai) and glomeruli (Aii) responses to repeated presentations of single-molecule odorants (1-Octanol *Oct*, leaf alcohol acetate *Laa*, 1-Hexanol *Hex*). Standard deviation projection across all stimuli (grayscale) of an exemplary animal, alongside false color-coded images of odor-induced responses, and the resulting overlay when treating each activation map as one channel in RGB color space. (B) Principal component analysis (PCA) of PN responses to five trials of each odor with an accompanying hierarchical binary cluster tree dendrogram (cophenetic correlation coefficient as *c*). Data correspond to (A). (C) Euclidean distance in PC space between responses of trials with the same odor (within) and different odors (between) of ten gregarious animals. Swarm plots show individual animal means, half-violin plots for probability density estimates, and crosses for grand means and 95% confidence intervals. (D) Confocal microscopy data of odor-induced responses to leaf alcohol acetate *Laa*, the locust odor *Lct*, and their combination *LvsLct*. Standard deviation projection across all stimuli (grayscale) of an exemplary gregarious animal, alongside false color-coded images after olfactory stimulation. (E) Average activity time courses of all gregarious animals (grand means with shaded 95% confidence intervals), swarm plots of individual animal means during the active period (time window indicated by dark gray bars), and crosses for grand means and 95% confidence intervals. (F) Stimulus interactions estimated using the Bliss score, distinguishing synergistic (score > zero) from antagonistic interactions between *Laa* and *Lct*. Swarm plots in Fi depict individual animal averages (split by antagonistic/synergistic regions) during the active period (E, dark gray bars), half-violin plots for probability density estimates, and crosses for grand means and 95% confidence intervals. The probability density function (PDF) in Fii shows the distribution of all Bliss score values between 15 for somata (red) and glomeruli (gray). (G-I) Correspond to D-F, but for solitarious animals. (J-O) Similar analysis as in D-I, but for recordings from the glomerular layer. Statistical inference in C, E, H, K, N was based on paired two-tailed bootstrap randomization tests. In all but C, *p*-value adjustments using the Bonferroni method. For both n.s.: not significant; p < 0.05: *; p < 0.01: **; p < 0.001: ***. Scale bars: 100 *µm*.

### 4.6 Confocal microscopy corroborates phase-dependent changes of antennal lobe activity

By using fine confocal optical slices (25 *µm*), we were able to separately analyze odor-response dynamics across different cell compartments. In both gregarious and solitarious locusts, we tested the responses to leaf alcohol acetate (*Laa*), the locust odor (*Lct*), and their combined mixture (*LaaLct*). We observed that each stimulus activates a broad population of PNs. This is exemplified by the response maps in Fig. 4D&G with the activity-based segmentation largely mirroring individual PN somata. Further corroborating our findings from the widefield imaging experiments above, the response magnitude of gregarious PN somata to the *LaaLct* mixture was significantly higher compared to the individual components *Laa* (*p* = 0.03) and *Lct* (*p <*< 0.001, paired two-tailed bootstrap randomization tests; Fig. 4D-E). Again, such a difference when adding *Lct* to *Laa* was not observed in solitarious animals (*p >* 0.99, paired two-tailed bootstrap randomization tests; Fig. 4G-H). This disparity can be partially attributed to an increase in the number of PNs recruited when presenting the *LaaLct* mixture in gregarious animals (24% vs 13% are *LaaLct*-specific in gregarious and solitarious locusts respectively; Tbl. 1). When recording odor responses from the PN dendrites (the glomerulus layer), we observed a congruent trend. In gregarious locusts, a larger proportion of glomeruli (19%) was *LaaLct*-specific compared to solitarious locusts (12%). Interestingly, the glomeruli response profiles were more transient in comparison to the sustained response patterns recorded from the somata. The average activity time courses of all active glomerular regions — medial tracts excluded — are shown in Fig. 4K&N. We also noted a minimal overlap between *Laa*-responsive and *Lct*-responsive glomeruli (gregarious: 3%, solitarious: 4%; Tbl. 1). The response overlap between *Laa* and *Lct* was much higher in the PN somata, as they integrate activity from multiple glomeruli and undergo additional processing with respect to PN dendrites.

Next, we summarised the odor response interactions of both phases for the mixture at the two cell compartments with a standardized measure that classifies pairwise interactions — Bliss score (Bliss (1939); Yeh et al. (2006)). The Bliss score distinguishes synergistic interactions (score > zero) from antagonistic ones (score < zero) between *Laa* and *Lct*, using Eqn. 3. When quantifying interaction strength by the average score across all antagonistic and synergistic regions for each locust, we observed a clear bias towards increased synergism in gregarious (Fig. 4FL) in comparison to solitarious locusts (Fig. 4IO). Although this trend was present in both somata (Fig. 4FI) and glomeruli (Fig. 4LO), the somata displayed a wider range of more extreme Bliss score values. This is evident in the overlaid distributions of Bliss interaction scores in Fig. 4F vs L and 4I vs O, highlighting the pronounced synergistic odor response interactions for the mixture in the somata.

### 4.7 Response motif dynamics in the antennal lobe reliably predict locust phenotypes

Our results repeatedly suggest that social cues synergize with appetitive olfactory signals in antennal lobe projection neurons of gregarious locusts, with the strongest interactions observed in the PN somata. In view of that, we now seek to characterize individual PN response profiles to uncover what distinguishes gregarious and solitarious phenotypes.

We applied an unsupervised analysis to all PN somata response profiles resulting from the three stimuli (*Laa*, *Lct*, *LaaLct*), identifying seven distinct odor-induced response motifs (Fig. 5A&B). Ordered by peak magnitude and timing, these motifs categorize PN responses as non-responding (motif 1), inhibitory (below baseline motifs 2 and 3), delayed response (motif 4), weak excitatory phasic (motifs 5), strong excitatory phasic (motifs 6), and excitatory sustained (motif 7). Examining the proportion of PNs in each response category, which were averaged across animals for each stimulus (Fig. 5Ci), revealed key differences between gregarious and solitarious locusts. Notably, solitarious animals exhibited a higher prevalence of the delayed response (motif 4), while gregarious animals displayed a more pronounced excitatory sustained response (motif 7) when exposed to the mixture (*LaaLct*; Fig. 5Cii). The change in activity profiles can be best understood by examining the dynamics of motif transitions between all three stimuli and within single PNs (Fig. 5D). We tracked how PN response motif assignments change with odors (*i.e.*, pairwise transitions between *Laa* and *Lct*, *Laa* and *LaaLct*, or *LaaLct* and *Lct*) and constructed heatmaps to summarize the occurrence of all motif transitions. For example, the cell at column 4 → 7 and row *Laa* → *LaaLct* demonstrates the proportion of PNs that showed motif 4 (delayed response) in response to *Laa* but motif 7 (sustained response) to *LaaLct*. The maps show a highly dynamic code with many transitions not only between non-active to active states but also in temporal response profiles (Fig. 5E). We noted that, while some transitions are clearly prominent (*e.g.*, transitions to motif 4 in solitarious animals), the number of combinations and interactions make the interpretation complex. Since, however, these motif transition maps contain the details of how responses shift with stimulus and phenotype across the PN population, we tested whether they can reliably decode the corresponding animal’s phenotype (*i.e.*, gregarious or solitarious). For this, we examined the predictive strength of a linear discriminant analysis (LDA) classification model, trained on gregarious and solitarious motif transition maps (see Methods for details). This analysis revealed that antennal lobe response dynamics can reliably distinguish gregarious and solitarious phenotypes with 92% correct predictions (Fig. 5F).

**Figure 5:**
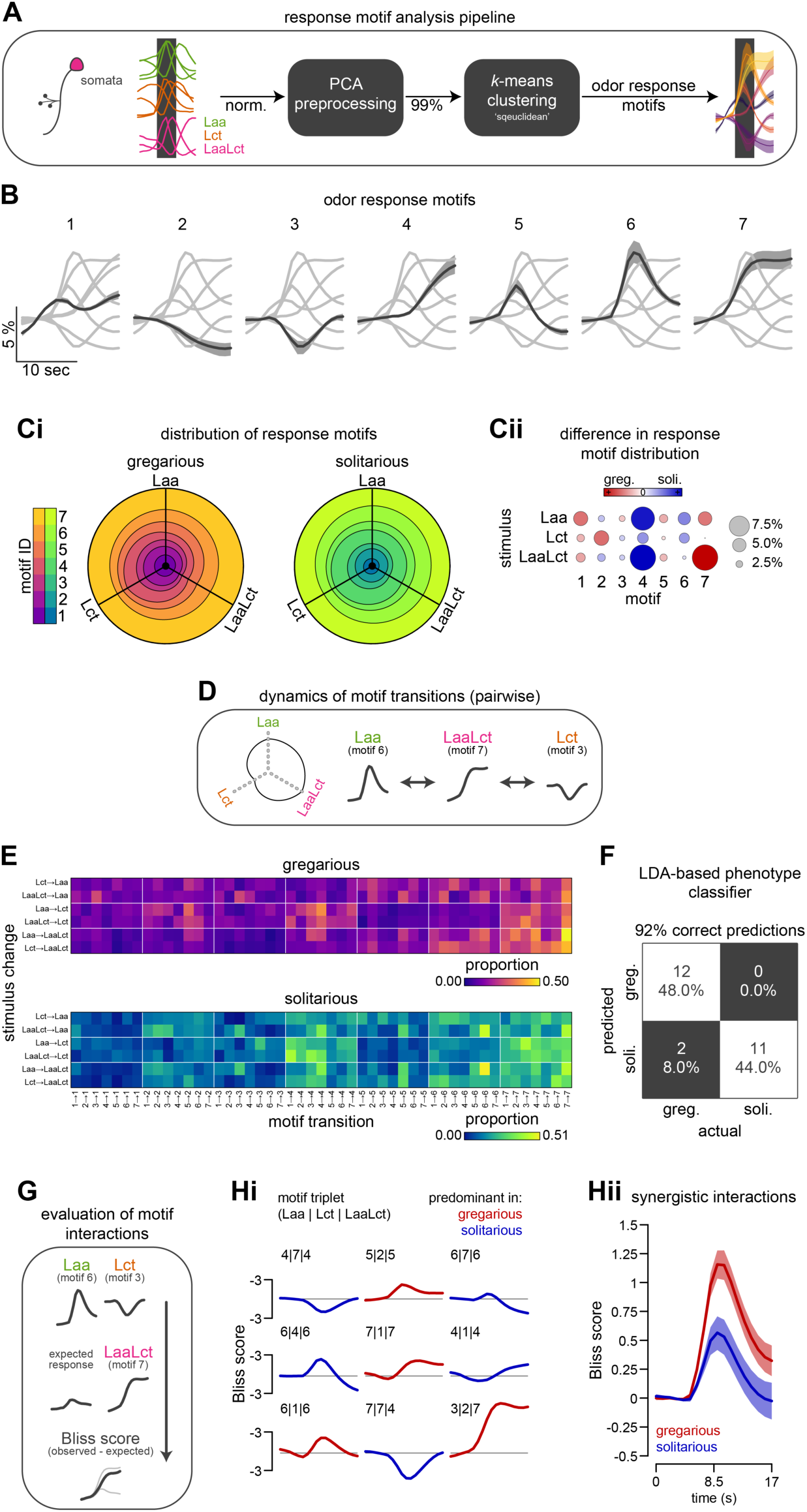
Analysis of dynamical odor response motifs. (A) Schematic illustration of the clustering process used to categorize the activity profiles of individual somata into seven distinct odor-induced response motifs. (B) Display of the centroids for each response motif, arranged according to peak magnitude and timing, shown as grand means across both gregarious and solitarious animals with shaded 95% confidence intervals. (C) Circular stacked bar plots represent the proportion of each response motif for each of the three stimuli (leaf alcohol acetate *Laa*, the locust odor *Lct*, and their combination *LvsLct*). Gregarious animals are shown on the left and solitarious animals on the right. Distinct differences between the two phases are emphasized in Cii, where variations in circle color and size denote the degree of disparity (absolute difference in proportion for each motif and odor). (D) Diagram demonstrating the dynamics of motif transitions, which lead to a unique motif triplet for every soma. (E) Visualization of the transition probabilities between response motifs, with color coding for the proportion of somata undergoing transitions between motifs (columns) across two stimulations (rows). Data for gregarious animals are in the top panel, with solitarious response transition probabilities below. (F) A confusion matrix showing correct (white) and incorrect (black) phenotype classifications using a linear discriminant analysis model. The model was trained on the dynamics of motif transitions for predicting each animal’s phenotype (see Methods for details). (G) Diagram showing the estimation of Bliss interaction score time courses of individual PNs based on their motif triplets. (H) Analysis of motif interactions using Bliss interaction score time courses to distinguish between synergistic (scores > zero) and antagonistic (scores < zero) interactions between *Laa* and *Lct*. Panel Hi shows the Bliss interaction score time courses for the nine triplet combinations that exhibited the largest differences in prevalence between gregarious and solitarious locusts (*cf.* suppl. Tbl. 1). Panel Hii displays the average (mean with shaded 95% confidence intervals) Bliss interaction score time courses based on all (but those with identical response motifs across all three stimuli) gregarious (red) and solitarious (blue) motif triplets (*cf.* D and G).

Particularly interested in the differences between gregarious and solitarious animals regarding mixture processing, we conducted a deeper analysis of the synergistic responses to the mixture (*cf.* Fig. 4). This analysis used the comprehensive data we have collected on the response motifs in individual PNs. We assigned each PN a response motif triplet, for instance, [6 |3| 7], which corresponds to the neuron’s response to the three presented stimuli (*Laa*, *Lct*, and *LaaLct*). Based on this information, we were able to calculate an individual interaction score (Bliss score; Fig. 5G). We observed that certain motif triplets, and thus certain Bliss interaction scores, were more prevalent in either gregarious or solitarious animals. We identified the nine triplets that most substantially differentiated the two phenotypes (Fig. 5Hi; color-coded according to the predominant phase) to further understand the crowding-induced synergy between food and social olfactory stimuli. Seven of these combinations involved PNs whose mixture responses were dictated by either the stronger (*e.g.*, 5 |2| 5) or the weaker component (*e.g.*, 4 |7| 4). However, the remaining two triplets (7 |7| 4 and 3 |2| 7, predominantly found in solitarious and gregarious locusts, respectively) displayed substantial mutual antagonistic and synergistic interactions (see bottom right of Fig. 5Hi). Pooling all the PN Bliss interaction scores, except those with identical response motifs for all three stimuli, highlights the differences in mixture interactions between the two phenotypes in the context of their response motifs (Fig. 5Hii). Taken together, our analysis suggests that network interaction variations are manifested in shifts in single PN response profiles, which conjointly give rise to a crowding-induced synergy in the olfactory processing of food and social odors.

**Table 1:** Average proportion (grand mean [95% confidence interval]) of granules responding in functional confocal laser scanning microscopy for the logical relationship between stimuli.

| Sets | somata (greg.) | glomeruli (greg.) | somata (soli.) | glomeruli (soli.) |
| --- | --- | --- | --- | --- |
| Laa only | 0.11 [0.08,0.14] | 0.12 [0.07,0.17] | 0.16 [0.10,0.21] | 0.15 [0.10,0.25] |
| Lct only | 0.12 [0.09,0.18] | 0.10 [0.07,0.13] | 0.14 [0.10,0.24] | 0.10 [0.07,0.14] |
| LaaLct only | 0.24 [0.15,0.36] | 0.19 [0.14,0.24] | 0.13 [0.09,0.20] | 0.12 [0.08,0.20] |
| Laa & Lct | 0.08 [0.05,0.12] | 0.03 [0.02,0.04] | 0.10 [0.07,0.13] | 0.04 [0.02,0.08] |
| Laa & LaaLct | 0.12 [0.08,0.16] | 0.14 [0.09,0.24] | 0.12 [0.09,0.16] | 0.08 [0.05,0.14] |
| Lct & LaaLct | 0.26 [0.17,0.39] | 0.40 [0.28,0.54] | 0.25 [0.19,0.32] | 0.46 [0.37,0.57] |
| all | 0.08 [0.05,0.12] | 0.03 [0.02,0.04] | 0.10 [0.07,0.13] | 0.04 [0.02,0.08] |

## 5 Discussion

In this study, we investigated the neural processing of olfactory information in changing social environments, allowing locusts to make decisions appropriately in a context-dependent manner. We established an *in vivo* functional calcium imaging protocol for reliably monitoring the activity of antennal lobe (AL) projection neurons (PN). Our results indicate a crowding-induced synergy in the processing of food and social odors that originates from phenotype-specific transition dynamics between distinct response motifs. As a direct consequence, this allowed us to use the AL network dynamics to reliably predict whether a locust was gregarious (group-living) or solitarious. Moreover, our study draws connections between the physiological observations and behaviorally tested patch preferences, providing further insight into the locust decision-making system during foraging.

During foraging, the presence of others can indicate both resource quality and location, offering graded and discrete information, respectively (Dall et al. (2005)). In gregarious locusts, this social information serves as an attractive cue for individuals to join a food patch, potentially even more than the sight/smell of the food itself (Fig. 1Ai and Günzel et al. (2023), also discussed in Lihoreau et al. (2017)). To estimate the relative contribution of the different cue types, we utilized a Bayesian decision-making rule inspired by Arganda et al. (2012). Our findings exemplify the universality of the Bayesian approach for disentangling contributing factors in behavioral data (as in Meyniel et al. (2015); Pérez-Escudero and de Polavieja (2011); Wystrach et al. (2015)), also when integrating more than two cue types. Our analysis indicates that patch selection is influenced by both visual and olfactory cues, and reveals phenotype-specific differences: while in solitarious locusts visual cues qualitatively predict patch choices on their own (similar preference in Vis(+)Olf(-) and Vis(+)Olf(+), Fig. 1C), it is only when both modalities are simultaneously present that a clear preference is demonstrated in gregarious locusts. Taken together, this revealed the nuanced roles different sensory modalities play in affecting locust foraging decisions, allowing for an investigation of the underlying neural mechanisms.

While it is known that the locusts’ phenotypic plasticity allows them to be well-suited for changing environments, the underlying neuronal mechanisms are not well understood. Building on the importance of olfaction for patch selection, we continued with mapping how food-related and social cues are being processed by the primary olfactory centers. Locust odor mapping has primarily been characterized as a dynamic population code, where the identity of an odor is determined by temporally evolving trajectories across an ensemble of PNs (Laurent and Davidowitz (1994); Laurent et al. (1996); Mazor and Laurent (2005); Stopfer et al. (2003)). Further studies strengthened the idea that this coding scheme is dynamic, history-, and context-dependent (Broome et al. (2006); Ling et al. (2023); Saha et al. (2015); Stopfer et al. (2003)). Pioneering *in vivo* functional calcium imaging in locusts to monitor the spatial arrangement of odor-evoked activity across the PN population in a single preparation, our study corroborates previous observations that individual odors activate broad sets of multiglomerular PNs. Similar to other insects, these subsets are odor-specific and stable across repetitions (Fig. 4A, in agreement with Mazor and Laurent (2005)). Furthermore, odor-evoked PN responses measured in different cellular compartments show substantial differences in temporal response properties: calcium responses in the glomeruli rise and decay fast with odor presentation. In contrast, responses in the somata are prolonged and temporally-varying (Fig. 4B&D vs. Fig. 4G&H, compatible with what is known about calcium influxes at the different compartments (Galizia and Kimmerle (2004); Lüdke et al. (2018)). To some degree, these differences can account for potential disparities between odor coding in the locust literature — described as temporally evolving trajectories (recorded electrophysiologically from PN somata, Anton et al. (2002); Mazor and Laurent (2005); Saha et al. (2015)), and the fruit fly and honeybee literature — describing more transient combinatorial patterns (recorded primarily via functional imaging from the dendritic terminals, *e.g.*, Sachse and Galizia (2003); Vosshall et al. (2000)). Recent observations from clonal raider ants — another insect species with largely arborized ALs, similar to locusts — revealed a highly stereotyped odor encoding logic in the ORN population (Hart et al. (2023)). As odor discrimination and coding efficiency are predicted to be preserved and even enhanced at the postsynaptic layer of PNs (Bhandawat et al. (2007); Lüdke et al. (2018); Raman et al. (2010)), it is likely that, at least, a comparable level of odor separation will be observed by the PN population. Taken together, these advocate for stable odor separation at the glomeruli across species and AL structures.

Our results repeatedly showed that PN responses to the leaf-associated volatile are modulated by phenotype (*i.e.*, with phase change) when mixed with the locust odor. In accordance with earlier studies showing no clear differences between the phases in AL anatomy (except for a higher number of sensilla in solitarious individuals, which may relate to their longer developmental period, Ochieng et al. (1998)) and in PN responses to single tested compounds (Anton et al. (2002); Ignell et al. (1998)), we have also observed a comparable response magnitude to the leaf and locust odors when presented separately. This suggests that phase change is less likely to impact presynaptic receptor sensitivity, and that the observed crowding-depended modulation is probably mediated by lateral network interactions. All other mixture responses, excluding the colony compound, were not higher than the response to their components, reflecting a hypo-additive, or elemental, mixture response (Clifford and Riffell (2013); Mohamed et al. (2019); Shen et al. (2013)). The comparison of PN response profiles to the leaf-locust mixture revealed higher mixture synergism in gregarious locusts, which is reflected by both larger proportion of mixture-specific PNs (Fig. 3C and Tbl. 1) and a larger number of sustained excitatory mixture responses (motif 7, Fig. 5Cii). The resulting odor profile, constructed from the maps of all response transitions, reliably separates gregarious and solitarious phenotypes (LDA analysis, Fig. 5), giving rise to specific differences in mixture processing. Most notably, we observed a higher proportion of PNs subjected to antagonistic mixture interactions in solitarious than in gregarious animals (Fig. 5Hi). Mixture responses dominated by the weaker component (*e.g.*, response triplet [4 7 4]), and ones with mutual suppression ([7 7 4]]) where more prevalent in solitarious locusts, while PNs subjected to synergistic mixture interactions were more prevalent in gregarious animals (*e.g.*, PN response triplets [5 2 5]] and [3 2 7]). Given the lack of observed excitatory local interneurons (LNs, Ernst et al. (1977)), these changes in network processing are likely mediated by disinhibition of lateral inhibitory LNs, innervated by *Laa*- or *Lct*-sensitive PNs. Local disinhibition will enable motif switches via the temporally diverse nature of LN populations (Nagel and Wilson (2016); Vogt et al. (2021)) and thus give rise to differential response profiles with increasing population density.

State-dependent regulation of odor processing in the insect antennal lobe has been predominantly investigated in the context of hunger, which incites animals to approach odor cues they would typically avoid (reviewed in Lin et al. (2019); Sayin et al. (2018)). This reconfiguration involves both modulation of presynaptic receptor sensitivity (Ko et al. (2015); Root et al. (2011)) and recruitment of local interneuronal input by neuromodulatory pathways (Albin et al. (2015); Vogt et al. (2021)). Another relevant type of context-dependent modulation in olfactory circuits is enhanced courtship receptivity in flies when they detect a food source nearby (Das et al. (2017); Grosjean et al. (2011)). This enhancement is mediated by lateral excitatory input to the pheromone-sensitive glomerulus in the presence of vinegar smell (Das et al. (2017)). Parallel to our observations, the context-dependent enhancement in Das et al. (2017) only occurs when both vinegar and the pheromone cVA are presented together, with PN responses to the single compounds remaining similar as in male and mated flies. It is therefore likely that the innervation pattern of local interneurons in the corresponding glomeruli may be sex-specific and potentially also context-specific (virgin vs. mated), thereby mediating differential lateral processing.

Swarms of desert locusts can span several hundred square kilometers and threaten the food security of millions. The density-dependent phase change from one phenotype to another is accompanied by substantial changes in the quantity, quality, and type of information animals can gather. Our results suggest that crowding already operates in early sensory processing centers, modulating — in our case — the antennal lobe output to influence animals’ foraging decisions. Yet, to fully understand locust phase change and swarm formation, future research is needed to address the multimodal nature of this phenomenon. Our Bayesian decision-making model suggests that the integration of olfactory with visual information is important for reliable decision-making, offering new research venues in the fields of vision and multimodal cue integration. Furthermore, our robust and data-driven calcium imaging analysis provides a comprehensive way to measure odor representation across the locust AL. Given the natural gradient of sociality in locusts, this can be further extended to investigate the cellular mechanisms of adaptive social behavior in changing sensory environments. Taken together, considering their behavioral and social plasticity, locusts not only provide an excellent opportunity for investigating olfactory processing in complex environments but also offer a trackable system for linking sensory adaptations to behaviorally relevant tasks.

## 6 Methods

### 6.1 Animals

All experiments were performed on gregarious and solitarious desert locusts *Schistocerca gregaria* (Forskål, 1775) obtained from our breeding colony at the animal facility of the University of Konstanz (Konstanz, Germany). Behavioral experiments were done with the last larval instar and functional imaging with freshly molted (< seven days old) adults. Gregarious locusts were reared in crowded cages (approx. 200 animals per 50×50×80 cm cage) while solitarious locusts were raised in individual boxes (9×9×14 cm) with opaque walls in a well-ventilated room with constant air exchange (following Hoste et al. (2002)). Except for animal density, all other conditions were kept similar for the two colonies with a 12:12h light:dark cycle, temperature of 27-29 °C, relative humidity of 45 % and a diet of in-house grown wheat seedlings, and freshly collected blackberry leaves (*Rubus* sect. *Rubus*). For all experiments, locusts of both sexes were used, after one night of starvation period.

### 6.2 Behavioural choice assay

We initiated our study with an analysis of the interactions between social and food cues in a patch selection assay (modified from Günzel et al. (2021)) which presented single locusts with four options: food (blackberry leaves, denoted as *Lvs*), a social setting (eight gregarious locusts, *Lct*), a combination of leaves and locusts (*LvsLct*), and an empty control patch (*Ctr*). Locusts were tested in a 90 cm diameter arena with the four patch choices distributed equally (exchanging positions between trials) in the arena, with a distance of approximately 7 cm from the wall. The four patch choices were made from identical circular plastic containers (width: 12 cm; height: 9 cm) and contained either 5 gr of ripped blackberry leaves, eight locusts, eight locusts with 5 gr of ripped blackberry leaves, or an empty control. In order to allow/prevent access to olfactory and/or visual cues, containers were either transparent with small holes, opaque with holes, or transparent but fully odor-sealed with Parafilm sealing film (Sigma-Aldrich, St. Louis, MO, USA). Each trial lasted 30 min and was recorded at 25 fps by a Basler camera (acA2040 - 90µm; Basler AG, Ahrensburg, Germany) with an IR longpass filter at the top of the arena. Uniform illumination was composed of a ceiling LED ring light (Hakutatz LED ring light, 33 cm, 35 W Bi-color 3200-5600 K) and a grid array of IR 850 nm with a diffuser plate below the arena. In order to prevent potential surrounding visual cues from the lab, uniform white walls of 1 m height were placed around the arena. Experiment containers as well as the arena were cleaned after each trial. The ambient temperature during experiments was kept between 24 and 26 °C.

### 6.3 Analysis of behavioral data

We obtained trajectories of individual locusts using a custom-written GUI (www.github.com/YannickGuenzel/BlobMaster3000) in MATLAB (R2021a, The MathWorks Inc, Natick, MA, USA) for automated tracking with manual supervision. Obtained trajectories were temporally smoothed by convolution with a Gaussian kernel (half-width = 2 s, 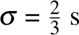). Visit density at each patch option was estimated as previously described (Günzel et al. (2021)) by binning *x/y*-coordinates in a 1000×1000 pixel grid, followed by a frame-wise application of a 2-D Gaussian smoothing kernel (s.d. = 78, spatial filter domain, filter size = grid size minus one; matching the density-independent interaction range employed by locusts of approx. 7 cm Buhl et al. (2006)). Resulting visit density maps were normalized by frame count and divided into four quarters, centered around each patch, and realigned such that trials could be pooled across different patch configurations.

To dissect the processes underlying locust patch choice, and to evaluate the contributions of the four information classes in the current assay (socio-visual, food-visual, socio-olfactory, and food-olfactory), we extended a Bayesian decision-making rule previously used in patch choice studies (Eqn. 1; Arganda et al. (2012); Günzel et al. (2023)). Given the availability of each information class (*e.g.*, *w_socio−ol f actory_*=1 for options containing locusts in the Olf(+) conditions, but 0 for the control option), we fitted the corresponding reliability parameter (*C_i_*) to the experimental visit density data in each patch quarter (*P_x_*).

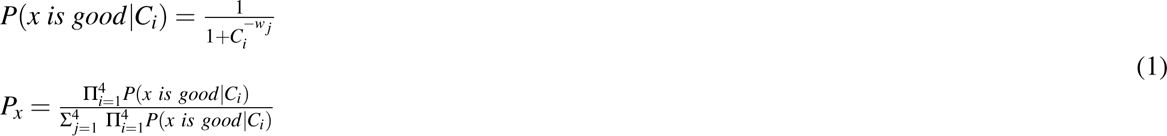

We used Bayesian hyperparameter optimization (MATLAB built-in function *bayesopt*) to fit the parameters in the range of 1*e^−^*^1^ and 1*e*^1^ (log-transformed). We repeated the fitting process 5000 times, resulting in a predictive distribution based on which we estimated 95% credible intervals. Moreover, the nature of parameter *C_i_* allowed us to estimate an information reliability score (*log*_10_(*C_i_*)) that assesses the degree of attraction (values approaching 1) or aversion (values approaching -1) each information class had on the tested locusts.

### 6.4 Preparation of animals for functional fluorescence microscopy

We performed widefield calcium imaging and functional confocal microscopy in the antennal lobes (ALs) of young adult desert locusts by modifying protocols for backfilling AL projection neurons with dextran-coupled calcium indicator (Paoli et al. (2017); Sachse and Galizia (2002)). For dye application, we anesthetized animals with *CO*_2_ to facilitate handling, removed wings and legs, and restrained the locusts in a custom-made holder using dental wax. On the first day, we opened a small window in the head capsule between the compound eyes, exposing the injection site. Next, using a glass capillary, we injected a crystal of a calcium indicator (Cal-520-Dextran Conjugate MW 10’000, AAT Bioquest | Biomol GmbH, Hamburg, Germany) into the medial calixes of both mushroom bodies for a retrograde backfill of AL projection neurons. Following the injection, the brain was covered with locust saline (Laurent and Davidowitz (1994)), and the head capsule was sealed with a drop of Eicosane (Sigma-Aldrich, St. Louis, MO, USA) to prevent brain desiccation. We allowed the dye to travel overnight with the animal kept in a humid box at 15 °C. On the following day, we exposed the brain and especially both antennal lobes. However, as the antennal base is located directly above the antennal lobes and consequently occluding them, we carefully relocated each antenna ventrally toward the ipsilateral compound eye. Special care was taken to not damage the antennal nerves. Following a successful relocation, we carefully removed the neural sheath covering the AL using fine forceps. For better optical accessibility, we gently elevated the brain with a stainless steel needle. As a last preparation step, we covered the preparation site with a transparent two-component silicon (Kwik-Sil, WPI, Sarasota, FL, USA).

### 6.5 Neuroanatomical staining

For morphological visualization of the olfactory pathway, we double-labeled PNs and ORNs using retrograde labeling. PN labeling followed a similar dye loading preparation as for the calcium imaging (see above) using the fluorescent tracer Alexa Fluor 568 conjugated with 10 kDa dextran. For receptor neuron tracing, the antennae were ablated close to the base, exposing the antennal nerve. The exposed antennal nerve was inserted into a glass capillary, loaded with 10 kDa dextran-conjugated Alexa Fluor 488 (both dyes by Thermo Fisher Scientific, Waltham, MA, USA). Animals were incubated for 24h at 15°C after which brains were quickly dissected and fixed in 4% paraformaldehyde (PFA) in phosphate-buffered saline (PBS) for 3h at 4°C in the dark. Brains were further washed in PBS and dehydrated in ascending concentrations of ethanol (50, 70, 90, 98, 100%), cleared in xylene for 10 minutes, and mounted in DPX mounting medium (Sigma-Aldrich, St Louis, MO, USA). Successfully stained antennal lobes were scanned as z-stacks with a Zeiss laser scanning microscope 880 equipped with an Airyscan fast detector and a 25x water immersion objective (LD LCI Plan-Apochromat 25x/0,8 Imm Korr DIC, Carl Zeiss, Jena, Germany) in the BioImaging facility at the University of Konstanz.

### 6.6 Widefield calcium imaging

We conducted calcium imaging analysis at a widefield fluorescence microscope (BX51WI, Olympus, Tokyo, Japan), equipped with a 20x water immersion objective (Olympus, UM Plan Fl 20x/0.50W), using an Omicron LED system, equipped with a 470 nm LED (Omicron-Laserage Laserprodukte GmbH, Rodgau-Dudenhofen, Germany) for light excitation with a 410 nm short-pass filter and a 410 nm dichroic mirror. In the emission pathway, the light was filtered through a 440 nm long-pass filter. Images with a spatial resolution of approx. 1.59 px/*µ*m were captured at 10 fps with a sCMOS camera (1024×1024 pixel with 2×2 on-chip binning; Prime BSI Express; Teledyne Photometrics, Tuscon AZ, USA).

### 6.7 Confocal laser scanning microscopy

Confocal microscopy of separate optical sections was acquired using an LSM 510 laser scanning confocal microscope (Carl Zeiss, Jena, Germany), equipped with a 20× water immersion objective (Zeiss W Plan-Apochromat 20×/1,0 DIC VIS-IR, Carl Zeiss, Jena, Germany). Sections were acquired at 1 Hz in two different depths (approx. 35 *µ*m, and 135 *µ*m from the surface of the AL), with a resolution of 0.77 *µ*m/px (*x*, *y*), and a full-width half maximum of 25 *µ*m (*z*).

### 6.8 Odor preparation, olfactory stimulation, and chemical analysis

For the widefield calcium imaging, odors were delivered via an automatic multi-sampler for gas chromatography (Combi PAL, CTC Analytics AG, Zwingen, Switzerland), which injected the odor pulse into a continuous flow of 1 mL/s of purified air with a matching injection speed. The stimulus odor was directed via a Teflon tube (inner diameter, 0.87 mm) to the antenna ipsilateral to the imaging site. For the confocal microscopy, odor stimulation was delivered via a custom-built olfactometer with a flow speed of 10 mL/s, and odors were injected for two seconds via a Teflon tube (inner diameter, 0.87 mm) to the ipsilateral antenna (Raiser et al. (2017)). Stimuli were delivered by activating valves that redirected air towards a vial containing 200 *µ*L of diluted odor. In the case of a single odor, the flow via the odor headspace (200 mL/min) was compensated by closing a balancer that reduces the airflow by the same amount. For mixtures, the headspaces of the two odors (100 mL/min each) were passed through a common Teflon tube, allowing them to mix before arriving at the locust antenna. A two-minute inter-stimulus interval of clean air was used to flush any residues of odors and to let the PNs return to their resting state. The flow of the individual odors and the mixtures was monitored by a photoionization detector (Mini-PID model 200A, Aurora Scientific Inc, Ontario, Canada), which was placed at the opening of the odor delivery tube.

All synthetic compounds (purchased from Sigma-Aldrich, Steinheim, Germany) were diluted in mineral oil. Locusts were tested with the following odorants: cis-3-Hexenyl-acetate diluted to 10*^−^*^2^ "*Laa*"), 1-Hexanol diluted to 10*^−^*^3^ "*Hex*" and 3-Octanol diluted to 10*^−^*^2^ "*Oct*"). Odour dilutions were chosen after an initial screening for average PN response magnitude comparable to that of the blackberry leaves *Lvs* in gregarious animals.

Lacking clear dominant candidates for emitted volatiles in the desert locusts (see latest summary of odor composition in Torto et al. (2021)), we used an extraction from our locust colonies as the social "*Lct*" odor. This was done by placing an evenly spread cotton wool layer for 2 days in a gregarious colony cage with nymphs of the last larval instar and immature adults. Leaf extract was produced by grinding blackberry leaves in mineral oil. Odor vials were filled with either 0.5 g of colony cotton wool, 0.3 g leaf extract, or 200 *µL* of odor dilution on 0.5 g of pure cotton wool. Mixtures were prepared by combining half the amount of the respective odor dilution, leaf extract, or colony cotton wool.

Volatile compounds for chemical analysis of the blackberry leaves were collected via 1h leaf extract exposure of solid phase microextraction (100 *µ*m Polydimethylsiloxane Coating SPME Fiber Assembly, Supelco, Bellefonte, USA) and analyzed with a TRACE GC Ultra Multi-channel gas chromatograph, coupled to DSQ II single quadrupole MS (Thermo Fisher Scientific, Waltham, MA, USA).

### 6.9 Analysis of calcium imaging data

For our calcium imaging analysis, we obtained a three-dimensional stack (x-y-time) for each stimulus and processed the data in MATLAB (R2022a, The MathWorks Inc, Natick, MA, USA). First, we filtered raw images spatially to account for shot (photon) noise, using a median filter (confocal microscopy: 3-by-3 pixels window; widefield imaging: 11-by-11 pixels window and frames were subsequently resized to 256-by-256 pixels using a box-shaped kernel). Next, we aligned frames belonging to one stimulus using the non-rigid motion correction algorithm NoRMCorre (Pnevmatikakis and Giovannucci (2017)). Then, stacks of the same stimulus set were registered based on phase differences between the stacks (Kughlin and Hines (1975)). Potential bleaching was accounted for by subtraction of a trendline (linear polynomial curve fit to the median pixel intensity time course of each stimulus). Carefully aligning our data allowed us to apply a box-shaped kernel (confocal microscopy: 3 frames; widefield imaging: 5 frames) in the temporal domain before calculating relative fluorescence changes 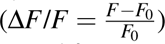 with the mean pre-stimulus activity before stimulus onset as baseline activity (*F*_0_; confocal microscopy: 5 frames; widefield imaging: 20 frames). In the case of two repetitions, pre-processed stimulus sets were registered based on phase differences between the sets and eventually averaged.

Interested in an unbiased evaluation of AL responses and detailed quantification of complex response patterns, we referred from manually applying circular regions of interest. Instead, we opted for segmentation into individual, activity-dependent regions, granules (for terminology *c.f.* solar granulation Blackwell et al. (1957)), based on the Voronoi topology of the standard-deviation-projection (std-projection) across the whole stimulus set. For this, we identified regional maxima and minima (pixel neighborhood was defined as pixels adjacent in the horizontal, vertical, or diagonal direction) in the standard deviation (s.d.) projection across all stimulus sets. Since the Voronoi diagram and Delaunay triangulation are geometric duals of each other, we could compute Voronoi granules from the Delaunay triangulation of the obtained regional extrema. This resulted in a rough 2-dimensional segmentation into regions that showed substantial changes in activity for any of the stimuli (regional maxima), separated by granules that lacked strong changes (regional minima).

We evaluated the activity of pixels along the Voronoi vertices to obtain a more refined segmentation. Specifically, we calculated the correlation coefficient between the activity of the pixel in question and the average activity of the neighboring Voronoi granule or the pixel’s ’home’ Voronoi granule, respectively. If the pixel was correlated more strongly with the neighboring Voronoi granule, we assigned it to this area in the next iteration. Alternatively, we kept the original assignment. This evaluation was applied to all pixels along borders between two granules and over a maximum of 100 iterations. Granules smaller than a minimum size (20 px in widefield imaging and confocal microscopy of somata; 5 px for confocal microscopy of glomeruli) were fused with their neighboring granules. Last, we removed regions lying outside the AL and classified all valid ones as either ’active’ or ’non-active’ given each stimulus. For this, we applied Otsu’s method to the mean intensity projection.

### 6.10 Construction of odor response vectors

To investigate network-spanning combinatorial coding patterns across all granules in the widefield imaging dataset, we created a response magnitude matrix including data from all gregarious and solitarious animals. In this matrix, each row represents a single granule, and each column corresponds to the respective mean response to each of the stimuli (*Laa*, *Lct*, *LaaLct*, *Hex*, *LaaHex*, *Oct*, and *LaaOct*). To account for inter-animal variability, we normalized the granules belonging to a single animal. For this, we initially subtracted the minimum value in order to root-transform the data. Next, we centered the animal’s data to have a mean of zero and scaled them to have a standard deviation of one. As a last step, we *z*-scored each granule separately before we projected the normalized response matrix into principal component (PC) space, retaining PCs that cumulatively explain at least 99% of the variance.

Following the preprocessing steps, we applied *k*-means clustering to construct distinct odor response vectors. The optimal number of clusters, *k* (1 *< k* ≤ 25), was ascertained by identifying the elbow point of Variance Ratio Criterion (VRC, Eqn. 2) curve, which was exponentially decaying with an increasing number of clusters. The VRC is defined as

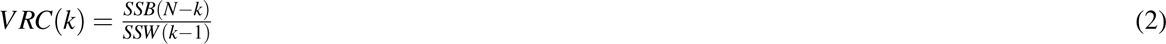

Here, *SSB* denotes the between-cluster variance, *SSW* denotes the within-cluster variance, and *N* denotes the number of observations, corresponding to the rows in the response matrix. We defined the elbow point as the point on the VRC curve that was farthest from the line segment connecting the first *VRC*(*k* = 2) and last *VRC*(*k* = 25) points on the curve. We performed the *k*-means clustering based on squared Euclidean distances, with a maximum number of 10^4^ iterations, 50 replicates, and with an online update phase in addition to the batch update phase, resulting in six odor response vectors.

### 6.11 Analysis of response consistency

To investigate the spatial representation of the general odorants used in our study (*Laa*, *Hex*, and *Oct*), we assessed the overlap in odor-induced responses. We performed functional confocal microscopy in the PN somata and glomeruli, repeating stimulus blocks (each stimulus presented once in a semi-random order) five times, to estimate the similarity between response maps. For this analysis, the normalized (range to [0, 1]) mean intensity maps from each animal were projected into the principal component (PC) space, treating each map as an observation and considering all pixels as features. We retained all PCs that cumulatively accounted for at least 95% of the variance. To evaluate the similarity between maps in PC space, we calculated the Euclidean distance between maps of the same odor stimulus and across maps from different stimuli. This allowed us to create a hierarchical cluster tree for each animal using the average linkage method. Further, we estimated the overall within-and between-stimulus Euclidean distances to pool results across all animals.

### 6.12 Analysis of synergistic stimulus interaction

To dissect odor response interactions, we analyzed them using the Bliss score (Eqn. 3), a standardized measure to categorize pairwise interactions (Bliss (1939); Yeh et al. (2006)).

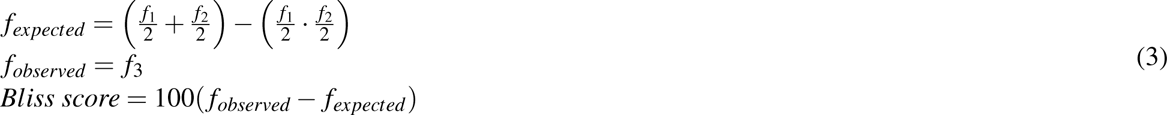

Here, *f*_1_ and *f*_2_ denote the individual response strengths (Δ*F*/*F* as decimal rather than percentage) to *Laa* and *Lct*, respectively, while *f*_3_ represents the observed response strength to the mixture (*LaaLct*). Positive Bliss scores, which arise when the observed response strength (*LaaLct* response) exceeds the expected response strength, indicate synergistic interaction effects between *Laa* and *Lct*. Conversely, negative values represent antagonistic effects, and values approximating zero indicate additive effects. Note that we divided *f*_1_ and *f*_2_ by two in Eqn. 3 to account for using only half of each component (*Laa* and *Lct*) when creating the mixture.

### 6.13 Analysis of dynamical odor response motifs for locust phenotype prediction

To gain deeper insight into the dynamics of odor-induced PN responses and their variation across different stimuli, we identified distinct temporal response motifs. For this, we adopted an approach comparable to the one used in the analysis of odor response vectors (see above). We used the PN somata confocal microscopy dataset, which encompasses data from all gregarious and solitarious animals, to construct a response profile matrix. Unlike the response vectors matrix, we treated all frames as features, and each soma appeared as three observations, one for each stimulus (*Laa*, *Lct*, and *LaaLct*). However, the normalization, principal component analysis-based preprocessing, and *k*-means clustering remained consistent, resulting in the identification of seven odor response motifs, based on the distinct patterns observed.

Next, we tracked how PN response motif assignments changed with odors. For this, we estimated the transition probabilities between response motifs (*e.g.*, motif 4 → motif 7) from one stimulus to another (*e.g.*, *Laa* → *LaaLct*) for each individual animal. We used this information to fit a discriminant analysis classifier to differentiate between gregarious and solitarious animals based on the dynamics of motif transitions(same diagonal covariance matrix for both phenotypes). The classifier was trained using a leave-one-out approach, with each iteration including all but one animal for training (the model’s costs and weights were adjusted to compensate for unequal sample sizes across classes) and employing the model to predict the phenotype of the excluded animal. This procedure was repeated for each animal in the dataset, providing a robust evaluation of the classification accuracy. Further, given the high feature dimensionality, we systematically reduced the feature set (transition probabilities), comparing different models using the Akaike Information Criterion to select the most appropriate model(Akaike (1973, 1974); Hurvich and Tsai (1993)).

### 6.14 Statistical analysis

All statistical analysis steps were conducted in MATLAB. Unless stated otherwise, we report averages as grand means, which represent the mean of animal means, together with bootstrapped 95% confidence intervals (with a bootstrap sample size of 5000) as shaded areas or error bars, respectively. Further, we used bootstrap randomization tests with *B* = 5 * 10^7^ samples (*cf.* MacKinnon (2009)) for statistical inference. We employed the following test statistic (Eqn. 4) for a paired, two-tailed comparison of observations *z* and *y* with the resulting pairwise difference *x* (mean 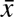, standard deviation *σ_x_*, sample size *n*)

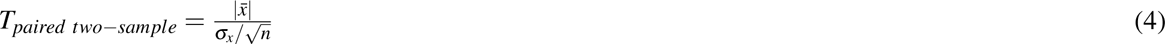

For an unpaired, two-sample, two-tailed test, comparing observations *z* (mean 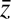, standard deviation *σ_z_*, sample size *n*) and *y* (mean 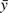, standard deviation *σ_y_*, sample size *m*), we used the following test statistic (Eqn. 5)

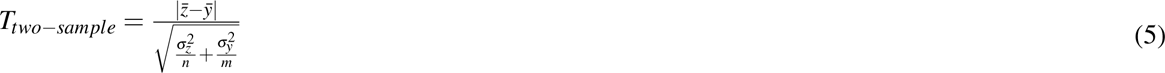

For both tests, we applied the same statistical summary to the permutated test statistic and to the test statistic of the original sample to determine the probability *p* of observing the permutated test statistic being the same or more extreme than the test statistic of the original sample. Note that this approach yields the lowest achievable *p*-value of 1*/B* = 2 *∗* 10*^−^*^8^. In the case of multiple comparisons, the *p*-value was adjusted using the Bonferroni method.

### 6.15 Data and code availability

All data presented in this paper and the custom code used for data analysis and figure creation will be publicly available upon publication of this manuscript.

## Material availability

This study did not generate any new reagents.

## Inclusion and diversity

We support inclusive, diverse, and equitable conduct of research.

## Author contributions

I.P, S.S, and E.C.F established the calcium imaging technique; Y.G and E.C.F conceptualized the project and wrote the first manuscript draft; I.P and S.K conducted behavior experiments, anatomy, chemical analysis, and functional imaging with S.S and Y.G providing conceptual and technical advice; Y.G developed analysis models, analyzed data and created all figures. All authors contributed to writing the final version.

## Acknowledgements

The authors thank Yvonne Hertenberger and Hannes Kübler for help with data collection; Nina Schwarz for providing locust illustrations; the team of the Bio Imaging Center at the University of Konstanz for help with obtaining confocal microscopy scans of antennal lobe double stainings; Marco Paoli for valuable insight on calcium imaging and corresponding analyses; Giovanni Galizia for providing feedback and facilities; James Foster, Ahmed El Hadi and Katrin Vogt for insightful comments on the manuscript. This work was completed with the support of the Deutsche Forschungsgemeinschaft (DFG, German Research Foundation) under Germany’s Excellence Strategy – EXC 2117 – 422037984.

## Supplementary information

### 1.1. Supporting figures

**Supplementary figure 1:**
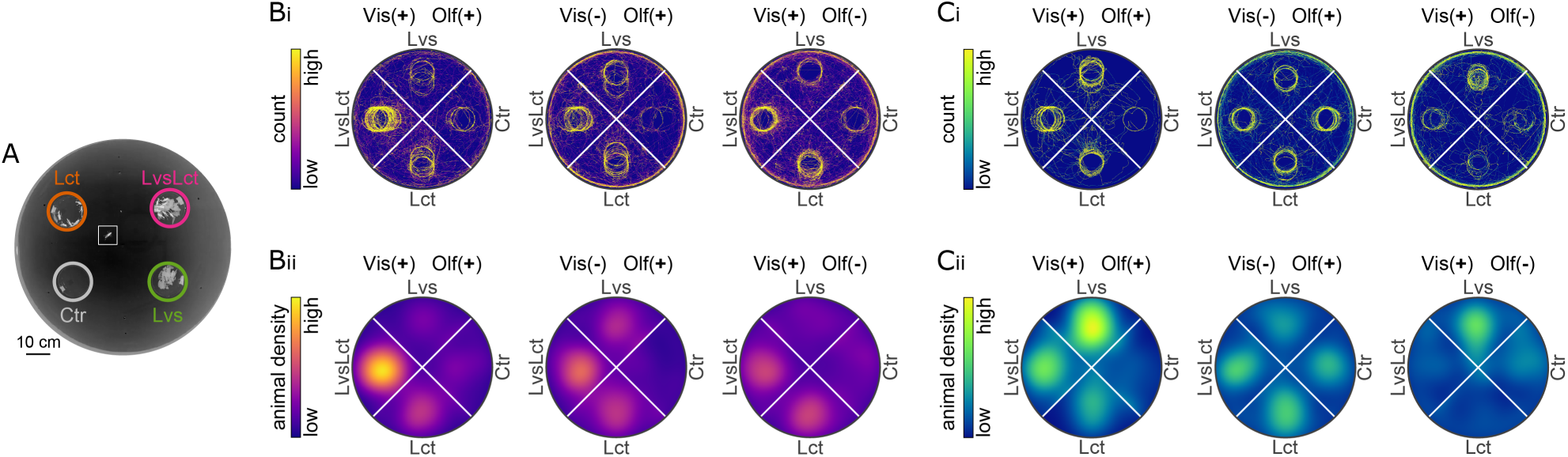
Details on patch selection experiments. (A) Animals were tested in a circular arena with four patch options (blackberry leaves *Lvs*, locusts *Lct*, leaves and locusts *LvsLct*, and control *Ctr*) under one out of three conditions (visual and olfactory cues available: *Vis*(+)*Ol f* (+); olfactory but no visual cues: *Vis*(*−*)*Ol f* (+); visual but no olfactory cues: *Vis*(+)*Ol f* (*−*)). Shown is a color-inverted, exemplary frame from testing a gregarious animal. The superimposed circles indicate the four patches, and the square shows the location of the locust. The animal was tested with both visual and olfactory cues available. (B) Bivariate histograms with 1000-by-1000 equally spaced bins were used to pool the 2-D trajectories of all tested animals (Bi), which in turn was used for estimating animal density in each stimulus quarter (Bii, same as in Fig. 1). (C) As in B, but for solitarious animals.

**Supplementary figure 2:**
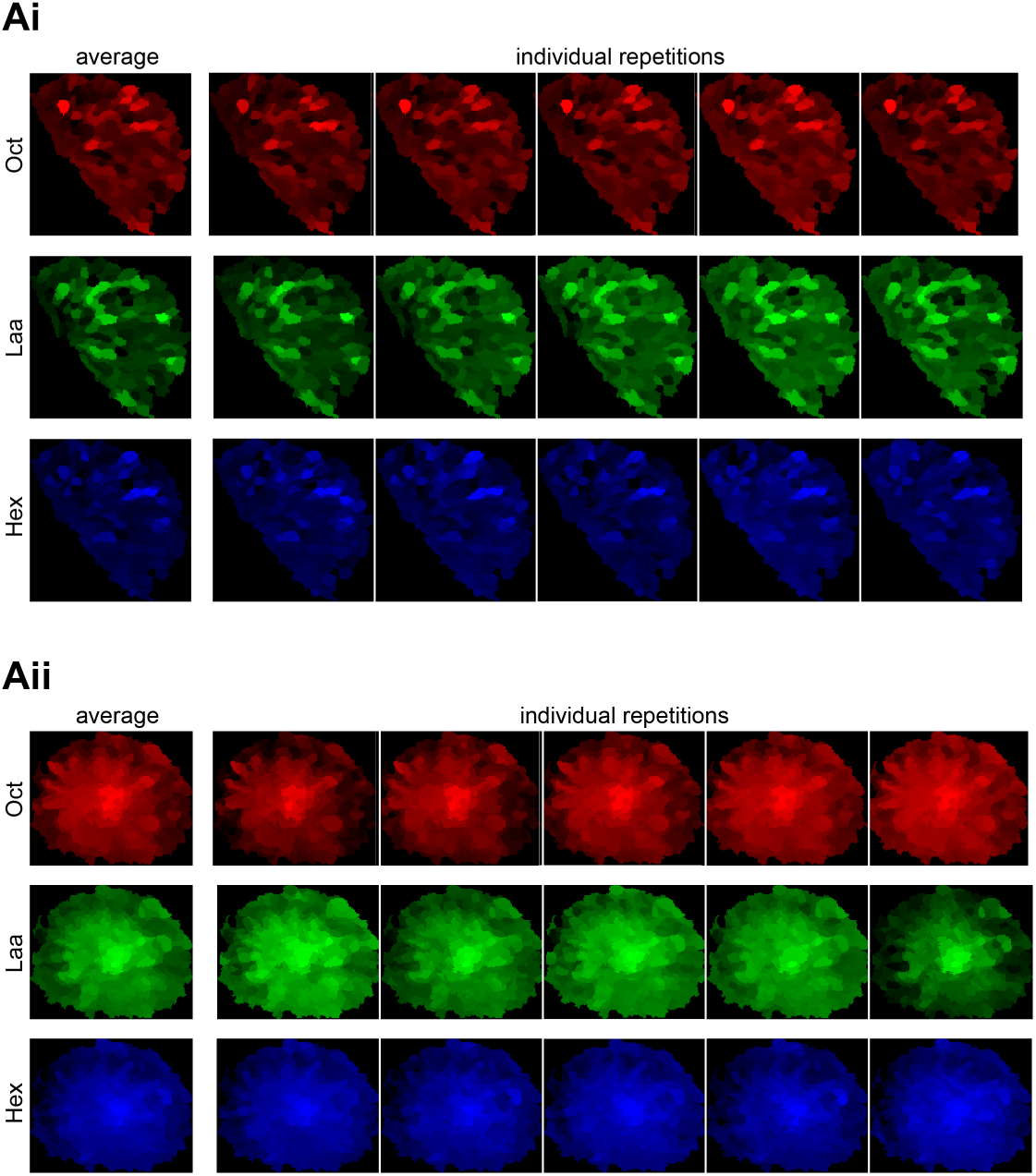
Details on response consistencies. (A) Individual trials of the example animal in Fig. 4A-B. Shown are odor-based color-coded mean intensity projections of projection neuron somata (Ai) and glomeruli (Aii) to repeated presentations of the odorants 1-Octanol *Oct*, leaf alcohol acetate *Laa*, and 1-Hexanol *Hex*. The left panels show average responses (as in Fig. 4) with the individual repetitions beside them.

### 1.2. Supporting tables

**Supplementary table 1:** Details on the prevalence of triplet combinations. We calculated how many times more a given motif triplet occurred than chance level (gregarious: 8.50; solitarious: 5.77) for both locust phenotypes. We report the nine triplet combinations with the largest absolute difference between gregarious and solitarious animals, matching the Bliss interaction score time courses in Fig. 5Hi. A value of 1 indicates that the triplet occurred as many times as expected by chance.

| motif triplet | greg. | sol. | abs. diff. |
| --- | --- | --- | --- |
| 4\|7\|4 | 1.65 | 11.78 | 10.14 |
| 5\|2\|5 | 7.06 | 1.56 | 5.50 |
| 6\|7\|6 | 3.88 | 8.84 | 4.95 |
| 6\|4\|5 | 3.53 | 7.80 | 4.27 |
| 7\|1\|7 | 4.94 | 1.04 | 3.90 |
| 4\|1\|4 | 0.35 | 4.16 | 3.81 |
| 6\|1\|6 | 5.53 | 1.91 | 3.63 |
| 7\|7\|4 | 2.24 | 5.72 | 3.48 |
| 3\|2\|7 | 3.30 | 0.00 | 3.30 |

